## Supplemental Figure 1 for "Reduction of HDAC2 expression in human induced pluripotent stem cell derived neurons improves neuronal maturation, mitochondrial dynamics and cellular neurodegenerative disease phenotypes"

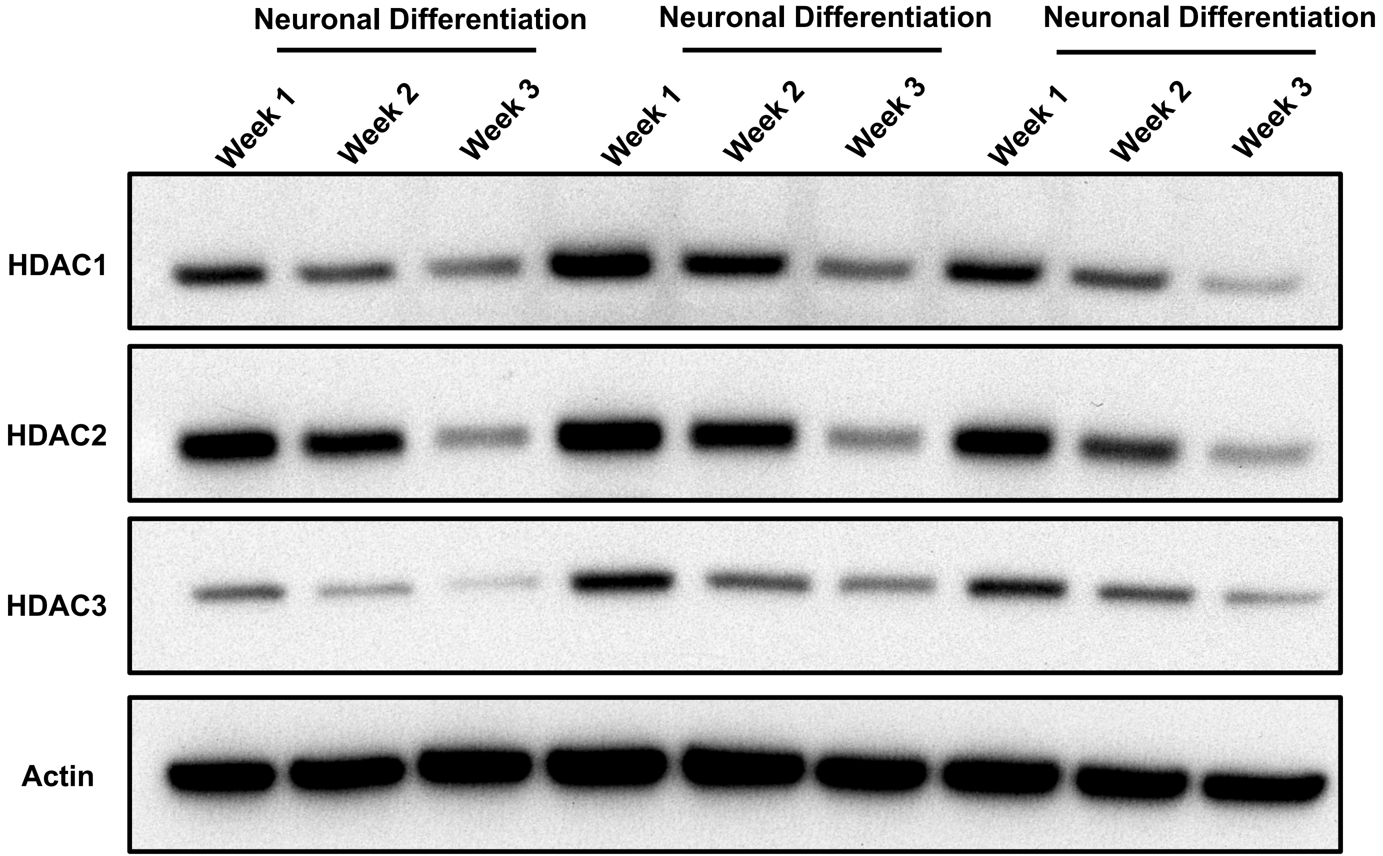


Supplemental Figure 1. Western blot of protein lysates from differentiatiating hiPSC-Ns shows that all Class I HDACs (HDAC1, HDAC2, HDAC3) decrease with neuronal differentiation
