## Supplementary figures and images for "Reduction of HDAC2 expression in human induced pluripotent stem cell derived neurons improves neuronal maturation, mitochondrial dynamics and cellular neurodegenerative disease phenotypes"

### Supplemental Figure 2

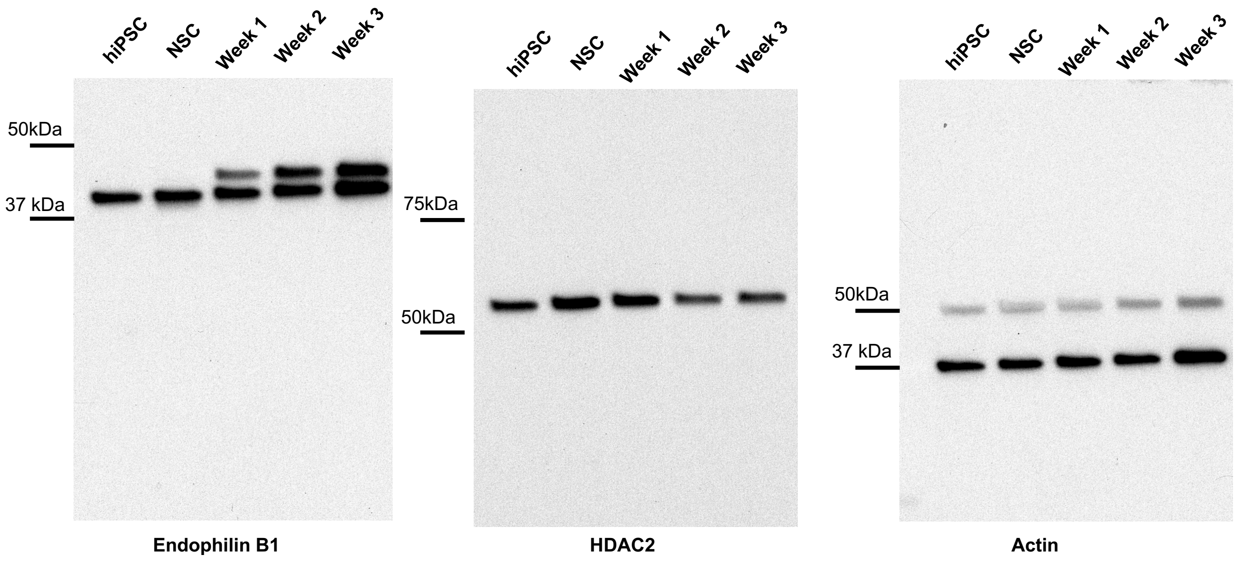


Western Blot: Fig 1

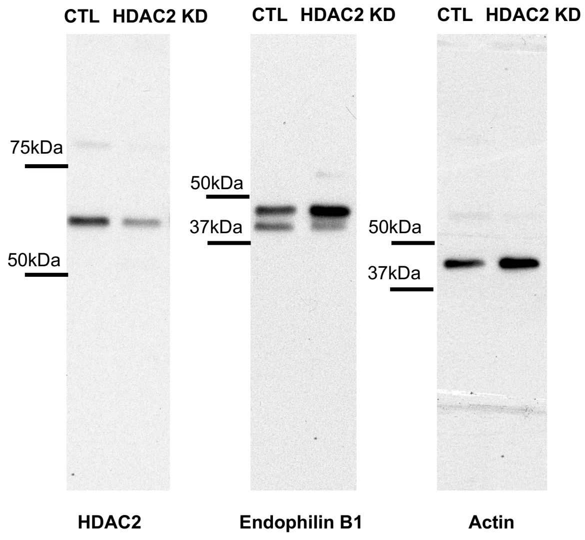


Western Blot: Fig 3

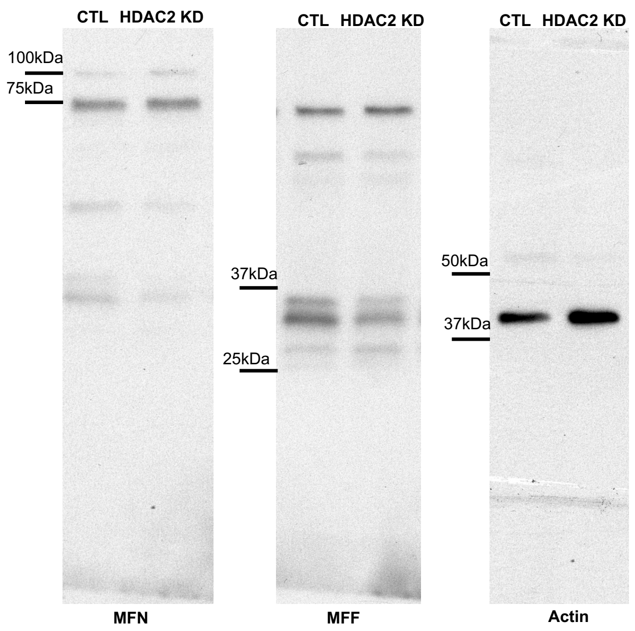


Western Blot: Fig 4

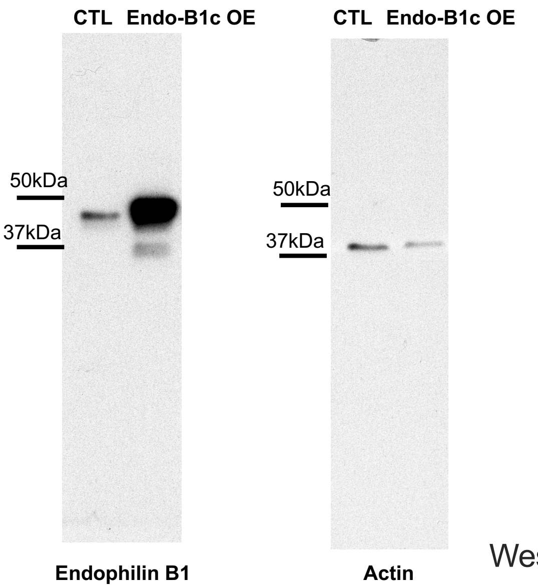


Western Blot: Fig 5

### Supplemental Figure 3

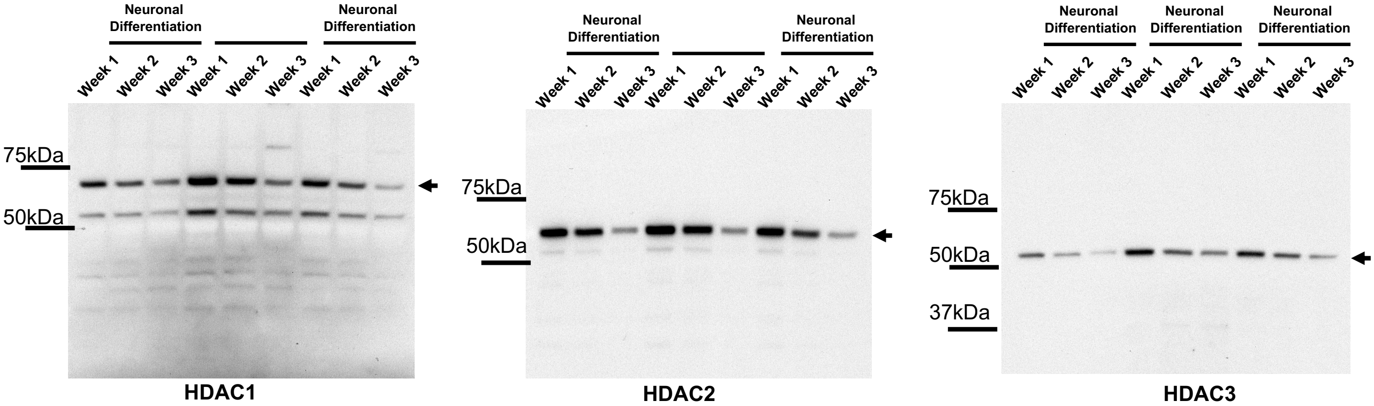

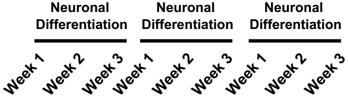

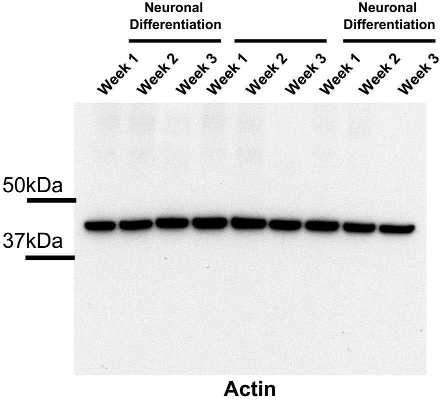


Western Blots for Supplemental Fig 1
